# Uncertainty in RNA-seq gene expression data

**DOI:** 10.1101/445601

**Authors:** Sonali Arora, Siobhan S. Pattwell, Eric C. Holland, Hamid Bolouri

**Affiliations:** Division of Human Biology, Fred Hutchinson Cancer Research Center, Seattle, WA 98109, USA

**Author notes:** Correspondence (E. C. Holland), (H. Bolouri).

## Abstract

RNA-sequencing data is widely used to identify disease biomarkers and therapeutic targets. Here, using data from five RNA-seq processing pipelines applied to 6,690 human tumor and normal tissues, we show that for >12% of protein-coding genes, in at least 1% of samples, current best-in-class RNA-seq processing pipelines differ in their abundance estimates by more than four-fold using *the same samples* and *the same set* of RNA-seq reads, raising clinical concern.

---

To explore the effects of RNA-seq data processing differences on gene expression estimates, we downloaded pan-tissue uniformly-processed RNA-seq abundance values from five different best-in-class processing pipelines (details in **Supplementary Table 1)** for 4,800 tumor samples from The Cancer Genome Atlas (TCGA), and calls from four pipelines for 1,890 normal-tissue samples from the Genotype-Tissue Expression (GTEx) project. We also included an additional dataset for the same samples with batch effect correction between TCGA and GTEx. To ensure fair comparisons, we limited all our analyses to protein coding genes that appear across all data sets (16,109 for TCGA; 16,518 for GTEx).

Differences among the RNA-seq pipelines discussed here include methods/software, software versions, run-time parameter values, and counting/normalization methods (**Supplementary Table 1**)^1^.

Several of the above data sources are only available in units of fragments per kilobase of transcript per million mapped reads (FPKM)^1^. However, FPKM counts do not take transcript lengths into account and are thus sensitive to differences in reference transcriptome annotations (**Supplementary Table 1**), as illustrated in **Fig. 1a,b**. To overcome this issue, we converted all data sets to units of transcripts per million (TPM) for a common set of genes^1^ (see **Online Methods**).

**Fig. 1.**
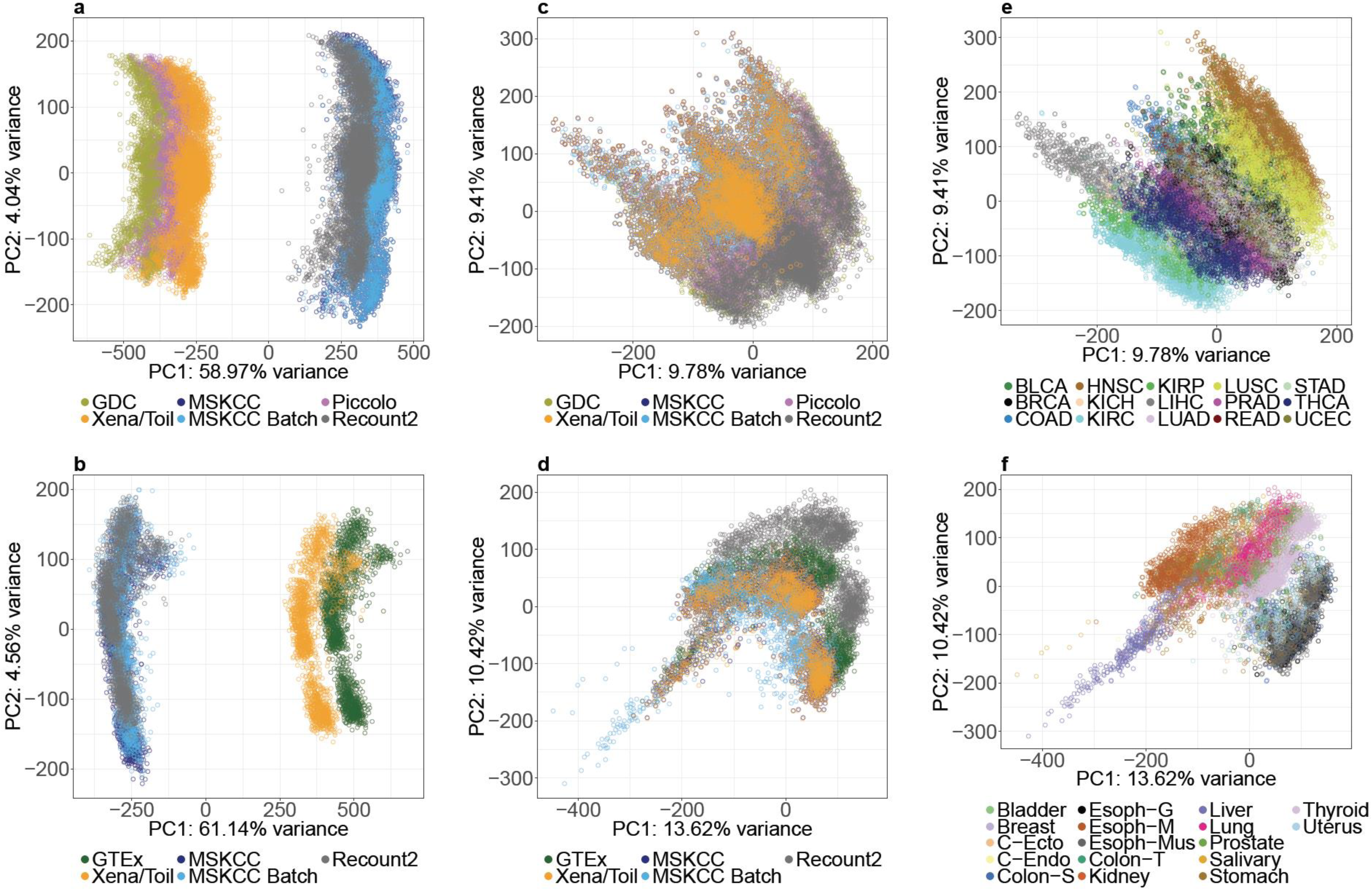
TCGA and GTEx data processed using diverse pipelines exhibit more variation by tissue source than by pipeline. (**a,b**) Principle Component Analysis (PCA) plots of TCGA and GTEx data using RPKM/FPKM values show very large inter-pipeline differences. (**c,d**) After transformation to TPM values, differences among pipelines are dramatically reduced. (**e, f**) Using TPM, variability between tissue types far exceeds inter-pipeline variability.

After the above unit conversion, the different versions of TCGA and GTEx data appear highly similar at the whole-genome level (**Fig. 1c,d and Supplementary Fig. 1,2**), and inter-tissue differences become more pronounced than data-source effects (**Fig. 1e,f)**. Moreover, inter-batch differences do not appear to confound comparisons *within* either TCGA or GTEx data sets, irrespective of data source (**Supplementary Fig. 3,4**), although comparisons *between* TCGA and GTEx samples may still be subject to batch effects^2^ (**Supplementary Fig. 5)**.

To quantify inter-pipeline differences not related to batch-effects, we analyzed data from the TCGA and GTEx projects separately and compared only pipelines that do not include batch-effect correction (five sources for TCGA; four sources for GTEx). We searched for genes whose expression in a given sample is >32 TPM according to one pipeline (suggesting high expression), but <8 TPM (i.e. >4-fold lower) in the same sample according to at least one other pipeline. To ensure that such large differences between abundance estimates are not due to a small number of outlier samples, we further required that the expression of a gene should be discordant (i.e. meet the above criteria) in at least 1% of all samples (48 samples for TCGA, 19 samples for GTEx). We refer to genes that meet all the preceding criteria as “discordantly quantified” genes.

We found 1,637 discordantly quantified genes (∼10%) in TCGA data and 1,214 discordantly quantified genes (∼7%) in GTEx data (**Fig. 2a, Supplementary Fig. 6)**. Across all TCGA and GTEx data sets, of 16,738 genes analyzed, 2,068 genes (12.36%) are discordantly quantified (**Supplementary Table 2a**). It should be noted that we imposed very stringent criteria in our identification of discordantly quantified genes. In particular, three and two-fold expression differences between pipelines affect many more genes (**Supplementary Tables 2b,c)**.

**Fig. 2.**
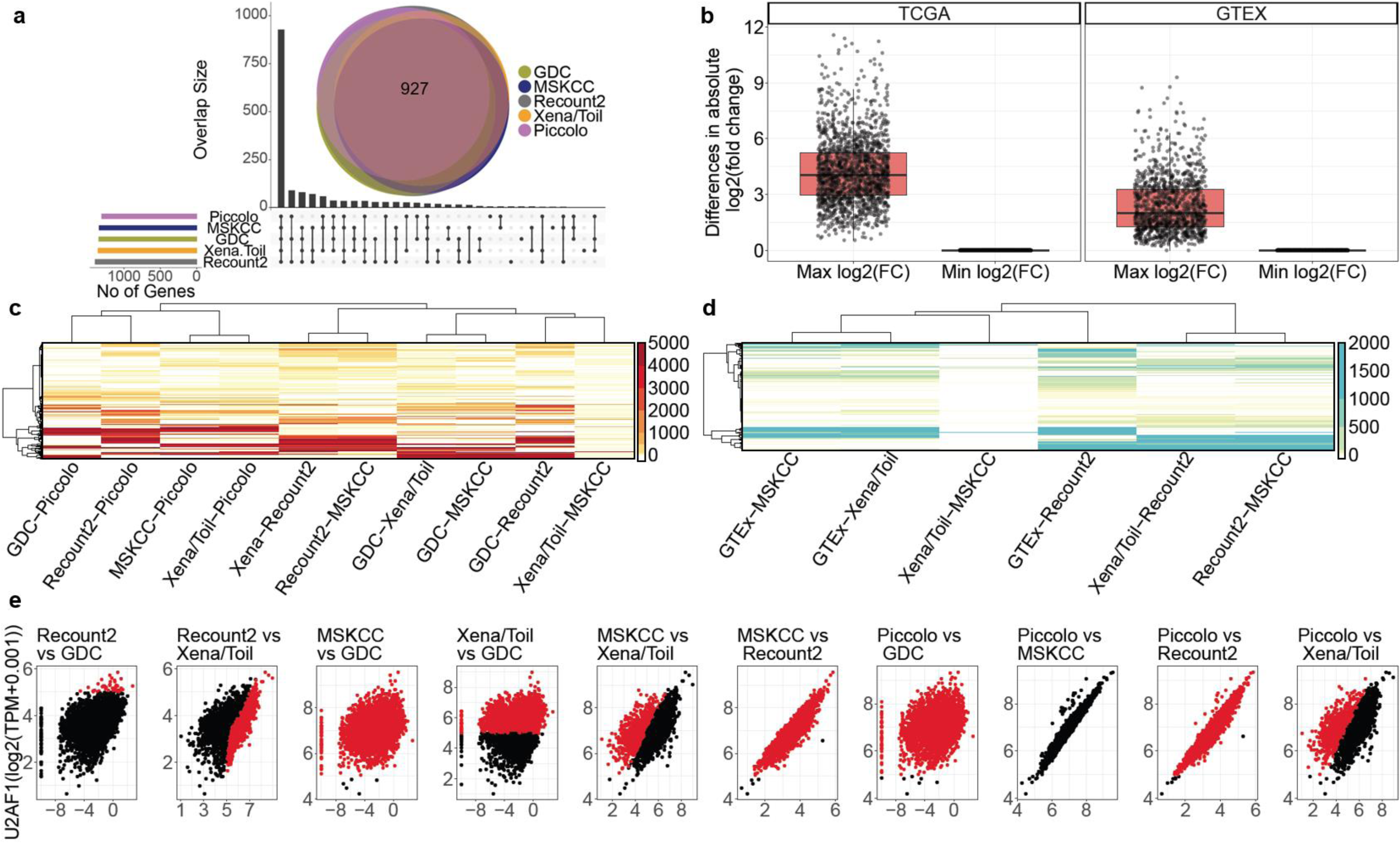
Large-scale inter-pipeline variability in specific genes. (**a**) Number of discordant genes per data source for TCGA, panel shows both UpSet^7^ and Venn diagrams. **(b)** Large inter-pipeline differences in fold change estimates for the same sample pairs among discordant genes. Shown are the maximum and minimum fold-difference estimates among pipelines for each discordant gene across all sample pairs. (**c, d**) Heatmaps showing the number of discordant samples per gene in TCGA and GTEx data. Each row represents one discordant gene, each column represents a comparison between two pipelines (as labeled). (**e**) Example pairwise comparisons of expression abundance estimates for U2AF1 showing diverse modes of disagreement among different pipelines. Panel titles specify the pipelines as Y-axis vs. X-axis.

The *relative* expression of a given gene between two samples may be expected to be more comparable across pipelines^3^. However, of the above 2,068 discordantly quantified genes, 1,958 genes (∼ 95%) have >2-fold inter-pipeline differences in fold-change estimates for the same sample pairs **(Fig. 2b and Supplementary Table 3)**

Our findings are consistent with those of the Sequencing Quality Control Consortium, which found that absolute RNA-seq abundance estimates were generally not trustworthy, and that reliable differential expression analysis was only feasible in less than two-thirds of the genome^3^. However, whereas SEQC compared expression data generated using multiple experimental and computational protocols, the discordant expression levels and fold differences reported here are for exactly the same RNA-seq reads, and arise entirely from differences in data processing pipelines.

Of note, many discordantly quantified genes have divergent expression values in large numbers of samples (>500, **Fig. 2c,d** and **Supplementary Table 4**). Importantly, the observed discrepancies are not attributable to a particular subset of processing pipelines or a particular subset of samples. Even the two pipelines with the greatest level of agreement (MSKCC and Xena/TOIL in the figure), still include multiple genes that are discordantly quantified in large numbers of samples.

For most discordantly quantified genes, differences between pipelines arise in a variety of ways. As an example, **Fig. 2e** shows the expression pattern of an example discordantly quantified gene (the splicing regulator U2 small nuclear RNA auxiliary factor 1 (U2AF1), which is frequently mutated in Myelodyplastic Syndrome^4^) in five versions of TCGA data (see **Supplementary Fig. 7a,b** for additional examples). We note that in some cases, estimates differ by a scaling factor. In other cases, U2AF1 is essentially not expressed according to one pipeline, but highly expressed according to another pipeline. In yet other cases, abundance estimates from two pipelines are simply poorly correlated. These large uncertainties in abundance estimates pose a significant challenge to biomedical research.

For example, more complex genome annotations can increase the numbers of unmapped and multi-mapped reads^5^. Indeed, 240 of our 2,068 discordantly quantified genes (11.61%) are known to be frequently affected by multi-mapping reads (**Supplementary Table 2**)^6^. Our list of 2,068 discordantly quantified genes includes 784 disease-associated genes (**Supplementary Table 5**), such as CEBPA, HIF1A, and KRAS, with important clinical implications.

In addition to poor inter-pipeline correlations in mRNA abundance (**Supplementary Fig. 8a,b**), for approximately half of the TCGA genes with available protein abundance data, mRNA and protein levels show remarkably low levels of correlation (**Supplementary Fig. 8c, Supplementary Tables 6a,b,c**) raising further concerns regarding the use of RNA-seq data for biomarker and target discovery.

The bioinformatics pipelines compared here represent best-in-class efforts by leading research teams, and utilize well-established, widely used methods. The differences among these pipelines arise from diverse implementation choices including statistical and algorithmic methods, software versions, and run-time parameters.

The discordant abundance and fold-change estimates revealed here do not imply any technical errors. Rather they highlight inherent uncertainty in processing noisy and complex data. Nonetheless, the end result is that for the discordantly quantified genes reported here (**Supplementary Table 2a**), we can have little confidence in the abundance estimates produced by any RNA-seq processing pipeline. For critical applications such as biomedical research and clinical practice, a concerted, community-wide effort will be needed to develop gold-standards for estimating the mRNA abundance of these genes.

## Acknowledgements

We thank Mary Goldman, Gavin Ha, Abhinav Nellore, Stephen Piccolo, Nikolaus Schultz, John Vivian, and Mark Ziemann for their comments on an earlier version of this manuscript. We thank Michael Gutteridge for help with Amazon S3.

## Author Contributions

All analyses were performed by SA and HB and discussed with SSP and ECH. All authors contributed to the final manuscript.

## Data and materials availability

**all data and scripts used in this manuscript are freely available via: https://github.com/sonali-bioc/UncertaintyRNA.**

## Online Methods

All analyses were performed in R (https://www.r-project.org/) using Bioconductor (https://www.bioconductor.org/) packages. To maximize transparency and reproducibility, we have deposited all scripts, associated data, and a large number of additional plots in a Github repository: https://github.com/sonali-bioc/UncertaintyRNA.

### Data sources

RNA Seq gene expression data was downloaded from five and four different sources for TCGA and GTEx respectively (Supplemental Table 1). Each source contained different numbers of genes and samples, thus we included only shared protein coding genes and samples found in every source in our analysis. The batch information for Sequencing Center, Tissue Source Site (TSS) and Plate ID were downloaded from https://gdc.cancer.gov/resources-tcga-users/tcga-code-tables. The nucleic acid isolation batch, genotype and expression batch data for GTEx samples were downloaded from the GTEx website (https://gtexportal.org/home/datasets).

### Conversion of abundance estimates to Transcripts Per Million (TPM)

All data sources except Xena/Toil provided FPKM RNA Seq gene expression data. For consistency, we converted all FPKM gene expression data to TPM data using the formula in ^1^.

### Principle Component Analysis (PCA)

Principle Component values were generated using the R function prcomp() using all genes and visualized with R package ggplot2.

### Discordantly quantified samples

For a gene to be discordant, its expression in at least one data set should be more than 32 TPM (i.e. log2 TPM more than 5) and the log2 fold change should be more than 2 (i.e. >4-fold difference in expression).

### Discordant fold changes

For all sample pairs within each data source, expression fold changes were calculated for discordant genes (as defined above) and compared with fold change differences across other data sources.

### TCGA-GTEx batch effects comparison (Suppl. Fig. 5)

Four pipelines provide both TCGA and GTEx data. To visualize potential batch effects between GTEx and TCGA data, we applied Principal Component Analysis to expression data for Stomach/STAD, Liver/LIHC and Thyroid/THCA samples from each pipeline in a manner similar to ^2^. In addition to these within pipeline comparisons, we also compared the original versions of the TCGA GDC data and the GTEx (v6) data using the same approach.

### Correlations

Pearson correlation was calculated using rcorr() function from Hmisc R package to compare TPM data from five different sources of TCGA data with the original GDC data for each gene. For protein-mRNA correlations, PANCAN12 protein abundance data was downloaded from https://xenabrowser.net/datapages/?dataset=TCGA.PANCAN12.sampleMap/RPPA_RBN&host=https://tcga.xenahubs.net. We calculated the Spearman correlation between protein levels and gene expression using rcorr() for only those genes for which we had both protein and gene expression data.

---

**Supplementary Figure 1.**
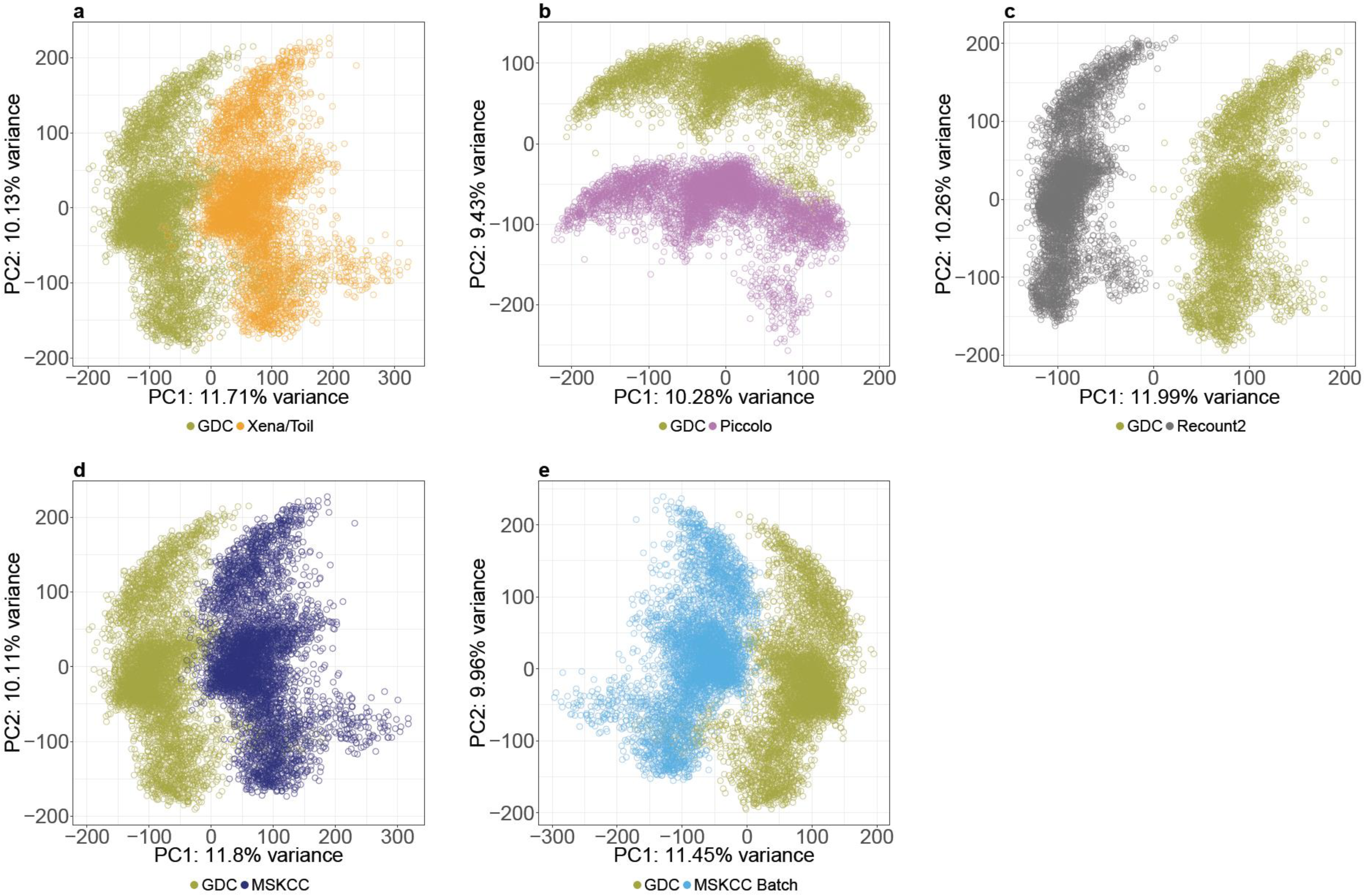
Principle Component Analysis (PCA) plots of TCGA data from **(a)** GDC and Xena/Toil, **(b)** GDC and Piccolo Lab, **(c)** GDC and Recount2, **(d)** GDC and normalized data from MSKCC (MSKCC) and **(e)** GDC and batch corrected data, after normalization from MSKCC (MSKCC Batch)

**Supplementary Figure 2.**
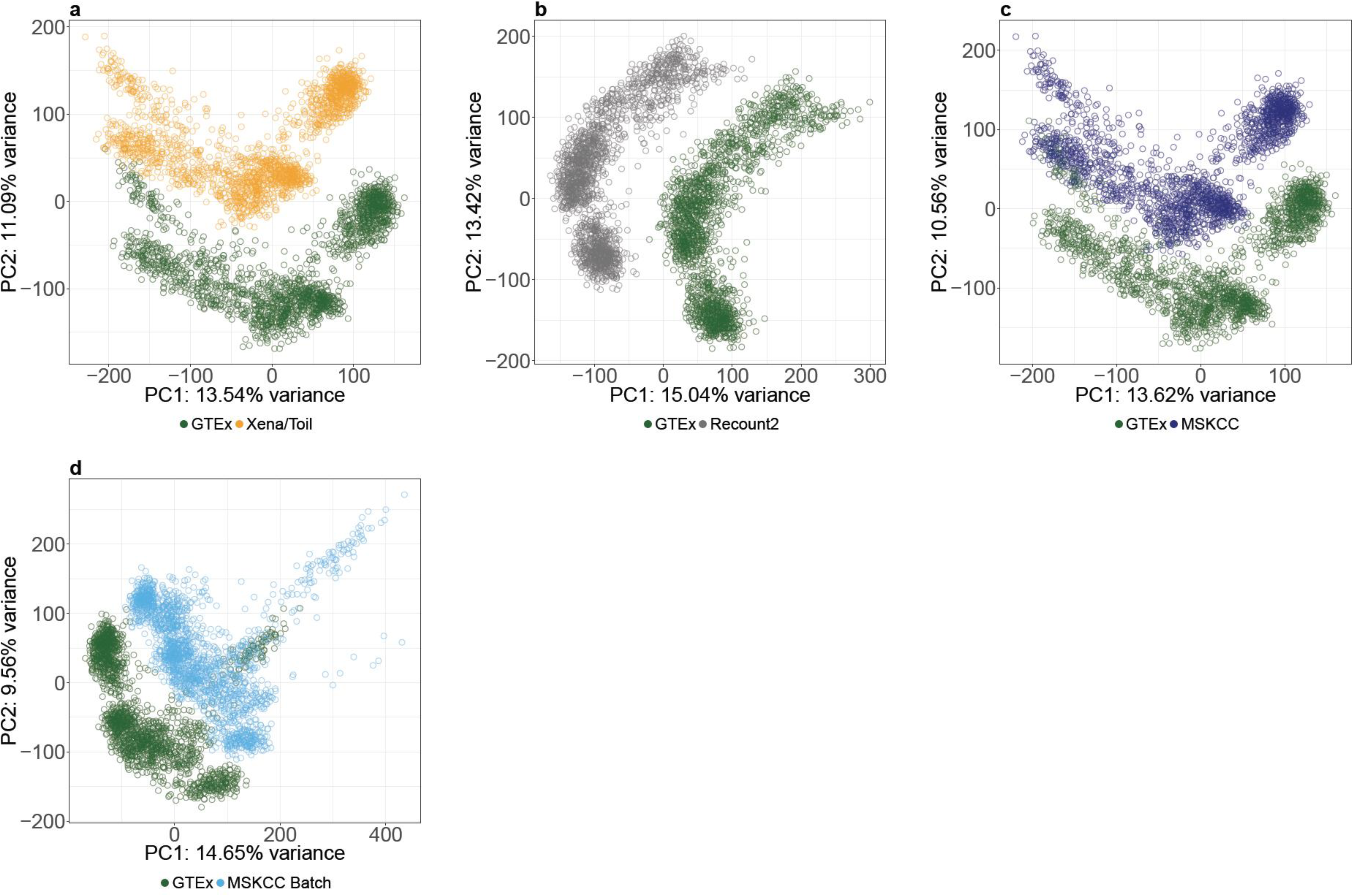
Principle Component Analysis (PCA) plots of GTEx data from **(a)** GTEx and Xena/Toil, **(b)** GTEx and Recount2, **(c)** GTEx and normalized data from MSKCC (MSKCC) and **(d)** GTEx and batch corrected data, after normalization from MSKCC (MSKCC Batch)

**Supplementary Figure 3.**
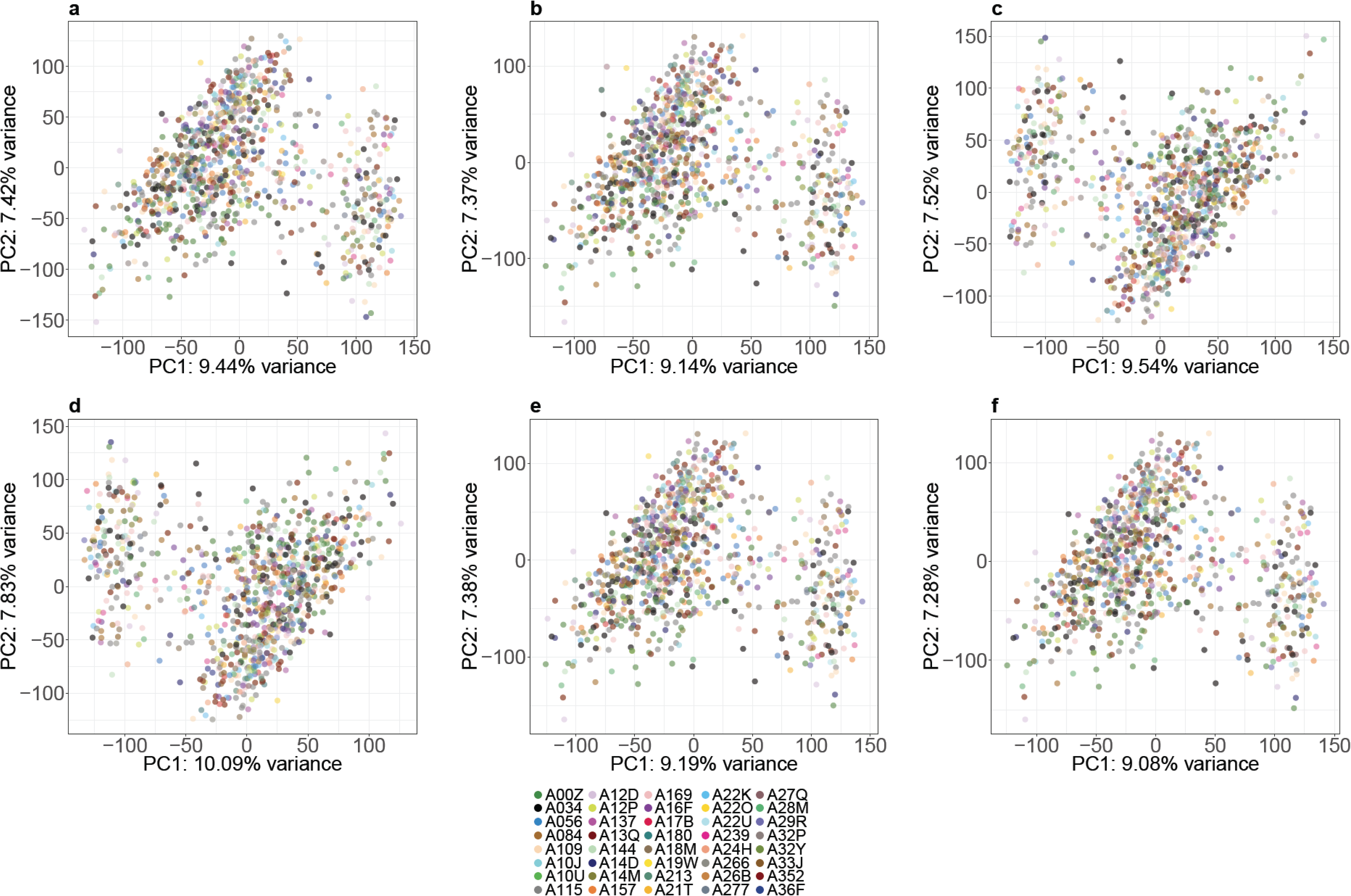
Principle Component Analysis (PCA) plots of TCGA data from **(a)** Recount2, **(e)** normalized data from MSKCC (MSKCC) and **(f)** batch corrected data, after normalizatio Batch) showing batch effects are not present. Colors in each PCA plot depict batches by “Plate Id”.

**Supplementary Figure 4.**
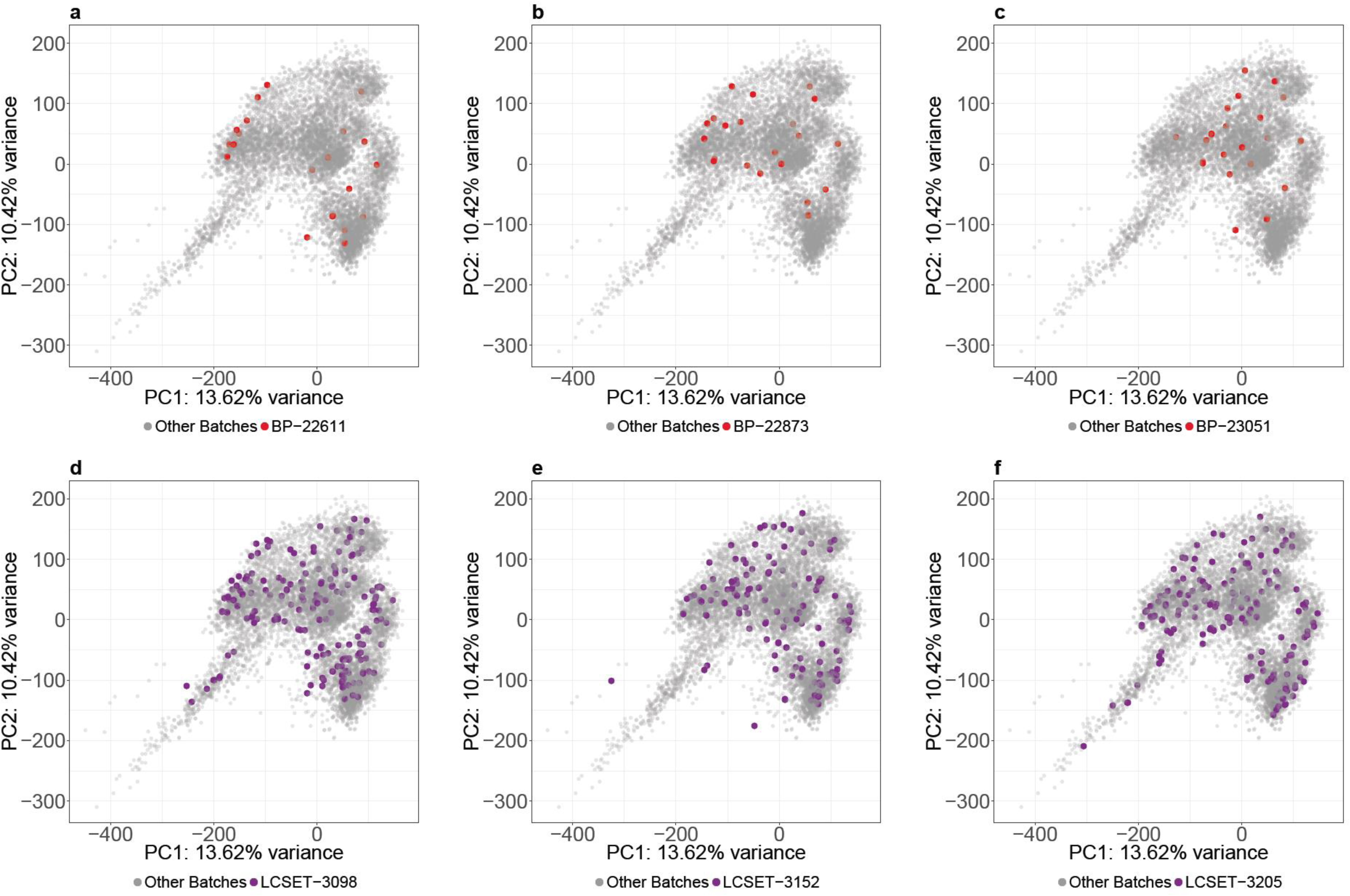
Uniformly processed GTEx RNA-seq data do not show batch effects. Two batch variables (nucleic acid isolation batch and genotype batch) are available for GTEx data. Principle Component Analysis (PCA) plots of GTEx data from all 5 sources of GTEx data is colored by three nucleic acid batches in red **(a)** BP-22611, **(b)** BP-23051, **(c)** BP-22873 and three genotype batches in purple **(d)** LCSET-3098 **(e)** LCSET-3152 and **(f)** LCSET-3205.

**Supplementary Figure 5.**
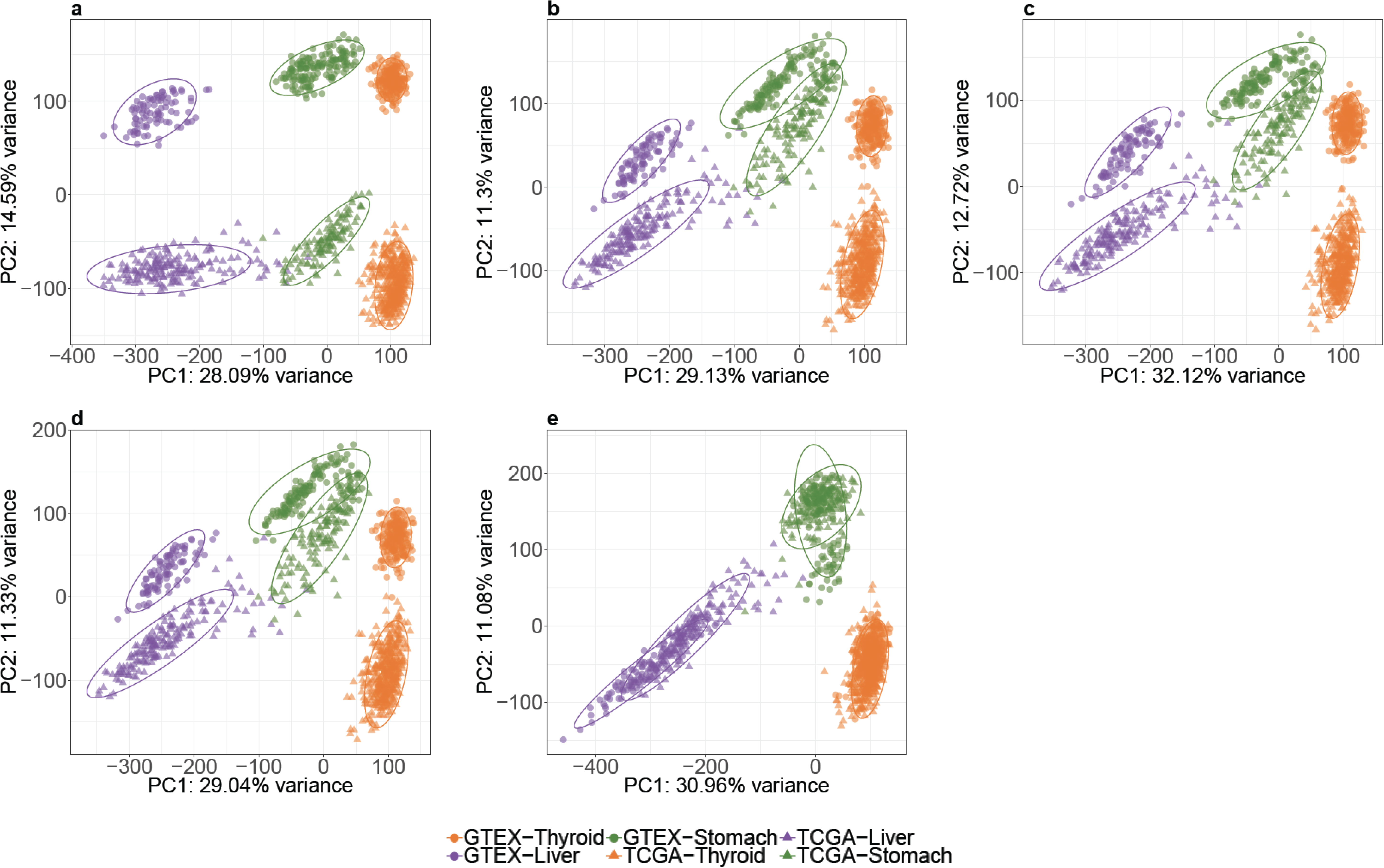
Illustration of potential batch effects between TCGA and GTEx. Gene expression data for thyroid, liver and stomach were compared with gene expression data for their respective cancer types (THCA, LIHC and STAD) from TCGA. **(a)** GDC, **(b)** Xena/Toil, **(c)** Recount2, **(d)** normalized data from MSKCC (MSKCC) and **(e)** batch corrected data, after normalization from MSKCC (MSKCC Batch) showing reduced differences between TCGA and GTEx.

**Supplementary Figure 6.**
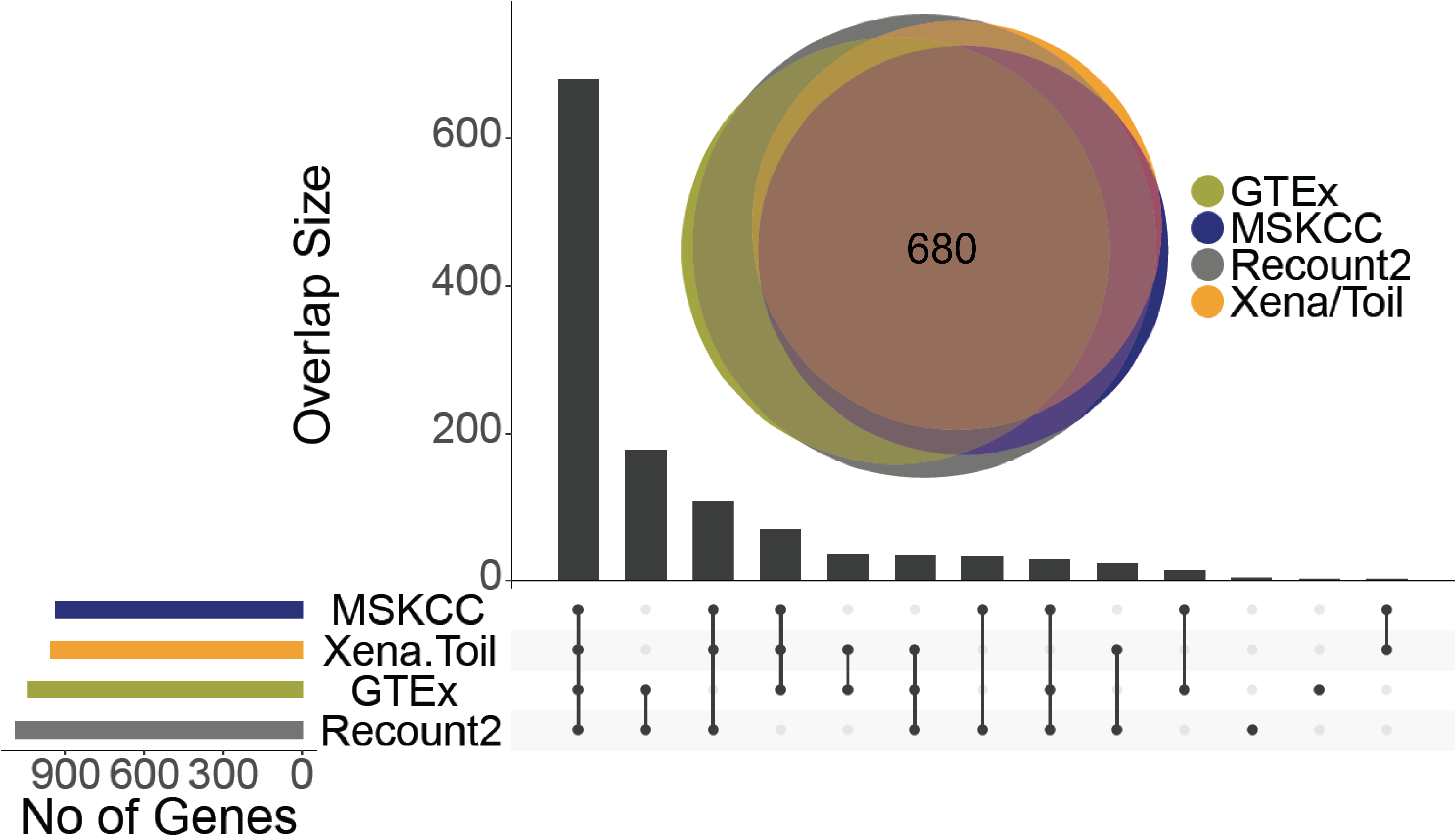
UpSet Plot and Venn diagram show number of discordant genes for GTEx data.

**Supplementary Figure 7.**
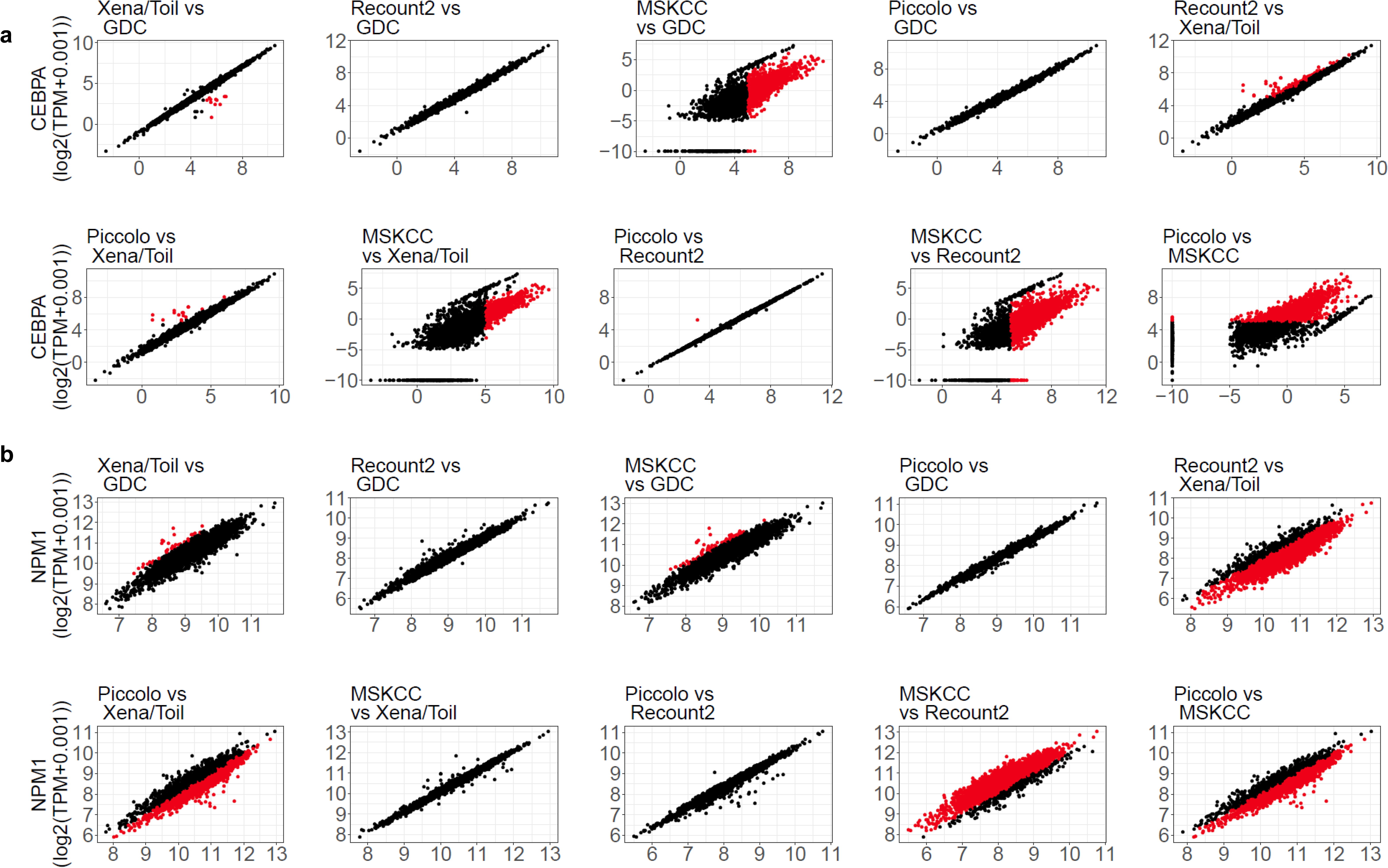
Pairwise scatter plots for gene expression [log2(TPM+0.001))] values for two genes CEBPA **(a)** and NPM1 **(b)**. Each panel is a comparison between two data sources, as indicated (figure titles show y-axis vs x-axis).

**Supplementary Figure 8.**
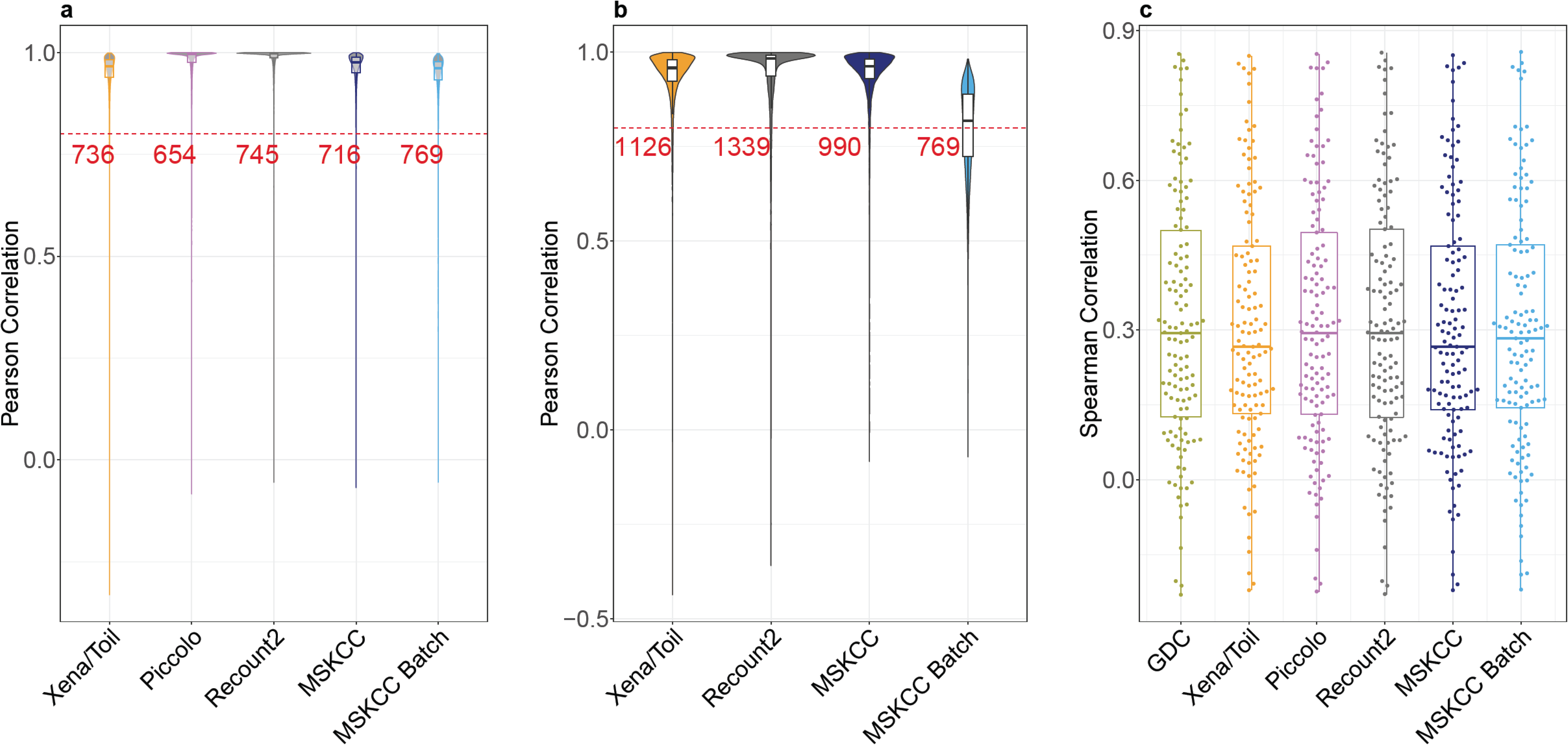
**(a)** Pairwise Pearson Correlation for gene expression data in TCGA samples comparing five uniformly processed data sources (as labeled) to data from the GDC (GDC-Xena/Toil, GDC-Piccolo, GDC-Recount2, GDC-MSKCC and GDC-MSKCC Batch). **(b)** Pairwise Pearson correlation for gene expression data in GTEx samples from four uniformly processed data sources compared to data from GTEx (GTEx-Xena/Toil, GTEx-Recount2, GTEx-MSKCC, GTEx Batch). **(c)** Spearman correlation for gene expression data from TCGA and available protein abundance data.

